# Growth performance and nutritional, mycochemical and bioactive profiling of *Volvariella volvacea* cultivated on tropical agricultural wastes

**DOI:** 10.64898/2026.07.30.741713

**Authors:** Olagunju Johnson Adetuwo

**Affiliations:** Department of Biological Sciences, Faculty of Science, Olusegun Agagu University of Science and Technology, Okitipupa, Ondo State, Nigeria

**Keywords:** Antioxidant, antimicrobial, bioconversion, cassava peel, Gmelina sawdust, oil palm pressed fruit fibre

## Abstract

Agricultural wastes such as Gmelina sawdust, cassava peels and oil palm pressed fibre are abundantly generated in tropical Nigeria and constitute environmental nuisance. This study evaluated the bioconversion potential of these wastes for cultivation of *Volvariella volvacea* and profiled the growth performance, nutritional, mycochemical and bioactive properties of the fruiting bodies. *Volvariella volvacea* was cultivated on three substrates: Gmelina sawdust (VVGS), oil palm fruit pressed fibre (VVOPF) and cassava peels (VVCP) in triplicates. Growth parameters including spawn run, pinhead formation, yield and biological efficiency (BE%) were determined. Molecular identification was confirmed using ITS, LSU and RPB2 markers. Proximate, mineral and vitamin compositions were determined by AOAC methods and HPLC. Mycochemicals were profiled by HPLC. Antioxidant activity was evaluated by DPPH, FRAP and total phenolic content, and antimicrobial activity by agar well diffusion and MIC against six clinical pathogens. All substrates supported mycelial growth and fructification with significant differences (p<0.05). Spawn run ranged from (10.1-12.0) days, with cassava peel best substrate showing the shortest period, oil palm fruit fibre showed highest BE among the three agro waste substrates of 74.2%. Fruiting bodies were rich in protein (10.2-12.3 mg/100 g/dry weight), high content of fiber, ash, and carbohydrate but low fat content. Abundant content of K, P, Ca, Fe and Zn. Vitamins B-complex and D2 were also detected. Phenols, terpenoid, alkaloids and saponins-like compounds were present with total phenols ranging from (1.5-1.63 mg GAE/g). Extracts exhibited dose-dependent DPPH scavenging (IC50 3.8-4.0 mg/mL), FRAP (10.10-10.2 mmol Fe2+ eq/g) and broad-spectrum antimicrobial activity (zones 11.9-17.4 mm, MIC 50.0 mg/mL), with cassava peel substrate showing highest bioactivity among the three agro waste substrates. Tropical agricultural wastes can be valorized for sustainable cultivation of nutritionally and bioactively rich *Volvariella volvacea,* with cassava peel and oil palm fruit fibre as most suitable substrate for food and nutraceutical applications in the study area.

## INTRODUCTION

The global demand for sustainable protein sources and functional foods has renewed interest in the cultivation of edible mushrooms through bioconversion of agricultural wastes (Adedokun *et al*., 2022; Chang & Wasser, 2017; Mayirnao *et al*., 2024). Edible mushrooms are low in fat and calories, rich in high-quality protein, dietary fibre, vitamins, minerals and bioactive mycochemicals, making them valuable for combating malnutrition and non-communicable diseases in developing countries (Barros et al., 2008; Reis *et al.,* 2012; Valverde *et al*., 2015; Rathore *et al*., 2017). Among tropical species, *Volvariella volvacea* Bull. ex Fr. Singer, commonly called paddy straw, oil palm or Chinese mushroom, is of significant economic importance due to its short cropping cycle of 10-14 days, ability to grow at high temperatures of 28-35 °C, and minimal requirement for complex cultivation technology (Chang & Miles, 2004; Stamets, 2000; Sánchez, 2010). The species is widely cultivated and consumed in tropical Asia and Africa and has been reported to possess antioxidant, antimicrobial, anti-inflammatory and immunomodulatory activities attributed to its phenolics, flavonoids, alkaloids and polysaccharides (Patel & Goyal, 2012; Ferreira et al., 2009; Wasser, 2014; Ferraro *etal,* 2021).

In Nigeria, large quantities of lignocellulosic agrowastes are generated annually from agriculture and agro-processing industries. Gmelina arborea sawdust is a major waste from sawmills in the rainforest zone, cassava peels constitute about 20-35% of cassava tuber weight from garri and starch processing, and oil palm fruit fibres are abundantly generated from palm oil mills in Southern Nigeria Adetuwo *et al*., 2026). These wastes are rich in cellulose, hemicellulose and lignin but are largely underutilized, often disposed of by open dumping or burning, leading to environmental pollution, greenhouse gas emissions and loss of potential resources. Bioconversion of these wastes through mushroom cultivation offers a sustainable, low-cost biotechnology for waste valorization, production of protein-rich food and generation of bioactive compounds in line with circular economy principles (Lavelli *et al,* 2018; Shirur *et al.,* 2021; Adedokun *et al*., 2022). The chemical composition of cultivation substrates significantly influences mycelial colonization, fructification, biological efficiency and the nutritional and functional quality of mushroom fruiting bodies (Chang & Miles, 2004; Ragunathan & Swaminathan, 2003; Sánchez, 2010). Substrates differ in carbon-to-nitrogen ratio, mineral content, lignocellulose degradability and presence of inhibitory compounds, which affect extracellular enzyme production by the mushroom and subsequent nutrient uptake (Chen *et al*., 2016; Silva *et al.*, 2002). Therefore, screening of locally available wastes is essential to identify optimal substrates that support both high yield and enhanced nutritional and bioactive quality of *V. volvacea* (Albertó, 2021). Previous studies have demonstrated successful cultivation of V. volvacea on paddy straw, cotton waste, banana leaves, sugarcane bagasse and oil palm empty fruit bunches (Chang & Miles, 2004; Sánchez, 2010; Stamets, 2000). However, information on the use of Gmelina sawdust, cassava peels and oil palm fruit fibres as sole substrates is very limited. More importantly, most existing studies focused primarily on agronomic parameters such as spawn run period, time to pinhead formation and yield, without comprehensive evaluation of the impact of substrate on detailed nutritional composition, mycochemical profile and bioactivities of the fruiting bodies (Reis *et al*., 2012; Ferraro *et al*., 2021). Nutritional studies that include proximate, mineral and vitamin contents, and functional studies covering total phenols, flavonoids, antioxidant and antimicrobial activities in relation to substrate type are scanty for these three wastes. Understanding substrate-induced variation in mycochemical composition and bioactivity is critical, as mushrooms grown on different lignocellulosic materials have been shown to exhibit significant differences in their antioxidant and antimicrobial potentials, which determine their value as functional foods and nutraceuticals (Ferreira *et al*., 2009; Reis *et al,* 2012; Effiong *et al*., 2024; Vamanu, 2012). The valorization of Gmelina sawdust, cassava peels and oil palm fruit fibres for *V. volvacea* production has not been comparatively evaluated under the same cultivation conditions in Nigeria, despite their abundance and disposal challenges.

Therefore, the aim of this study was to evaluate the growth performance of *Volvariella volvacea* cultivated on Gmelina sawdust, cassava peel and oil palm fruit fibre wastes and to profile the nutritional, mycochemical, antioxidant and antimicrobial properties of the fruiting bodies. The findings will provide insights into the bioconversion potential of these tropical wastes and their influence on the functional quality of *V. volvacea* for food and biotechnological applications.

## 2.0 MATERIALS AND METHODS

### 2.1 Study area and substrates

The study was conducted in Okitipupa (6.5025°N, 4.7795°E), Ondo State, Nigeria, characterized by tropical climate with annual rainfall 1,200-1,500 mm and temperature 26-34°C, favourable for *Volvariella volvacea* cultivation. Three agro-wastes abundant in the area - Gmelina arborea sawdust, oil palm pressed fruit fibre and cassava peels - were collected from sawmills and processing centres in Okitipupa, air-dried, milled and sieved.

Experimental design. Three experimental groups were set up for *V. volvacea* cultivation: VVGS (Gmelina sawdust), VVOPF (oil palm pressed fruit fibre) and VVCP (cassava peels). Each group was prepared in triplicate (1 kg dry substrate per replicate).

### 2.2 Substrate preparation and cultivation

Substrate moisture was adjusted to 60-65%, packed in 500 mL heat-resistant flat plastic containers and sterilized at 121°C for 1 hour. Grain spawn was prepared using sorghum grains supplemented with 2% CaCO_3_ and 1% gypsum, inoculated with pure *V. volvacea* culture and incubated at 28±2°C for 10-14 days until full colonization (Chang & Miles, 2004; Stamets, 2000; Sánchez, 2010).Each container was inoculated with 5-10 g spawn under aseptic conditions in laminar flow cabinet and incubated in darkness at 28±2°C, 75-85% RH for spawn run. Fruiting was induced at 25-28°C, 85-90% RH with 12 h diffused light and twice daily sprinkling. Mushrooms were harvested at cap expansion before spore release. Yield parameters including biological yield and biological efficiency [% BE = (fresh mushroom weight / dry substrate weight) x 100] were recorded.

### 2.3 Molecular profiling

Genomic DNA was extracted from fresh mycelium using CTAB method (White *et al*., 1990). Three markers were amplified: ITS (ITS1/ITS4), LSU (LR0R/LR7) and RPB2 (fRPB2-5F/fRPB2-7cR) using PCR conditions: 95°C 5 min, 35 cycles of 94°C 30 s, 55°C 45 s, 72°C 1 min, final extension 72°C 10 min. Products were verified on 1.5% agarose gel, Sanger sequenced, consensus sequences generated and confirmed via BLASTn (≥98% identity) in NCBI GenBank (Schoch et al., 2012; Adetuwo *et al*., 2026).

### 2.4 Nutritional Analysis

Freshly harvested fruiting bodies of *Volvariella volvacea* were cleaned to remove adhering substrate particles and other debris without washing to avoid nutrient loss. Samples were sliced into uniform pieces (approximately 3-5 mm thick) and oven-dried at 60 ± 2°C to constant weight (48-72 h). The dried samples were ground into a fine powder using a laboratory mill and passed through a 0.5-mm mesh sieve to obtain a homogeneous sample. The powdered samples were stored in airtight polyethylene containers at 4°C until analysis. All analyses were carried out in triplicate, and the results were expressed on a dry-weight basis unless otherwise stated.

#### 2.4.1 Proximate Composition

The proximate composition of the mushroom samples was determined according to the AOAC (2019) standard analytical procedures.

##### Moisture Content

Moisture content was determined by drying approximately 5 g of the fresh mushroom sample in a hot-air oven maintained at 105°C until a constant weight was obtained. Moisture content was calculated as the percentage weight loss during drying.

##### Crude Protein

Crude protein was determined using the Kjeldahl method (AOAC 984.13). Approximately 0.5 g of dried mushroom powder was digested with concentrated sulfuric acid in the presence of a catalyst mixture until a clear digest was obtained. The digest was neutralized with sodium hydroxide, distilled, and the liberated ammonia was collected in boric acid solution before titration with standardized hydrochloric acid. Total nitrogen was multiplied by a conversion factor of 6.25 to estimate crude protein content.

##### Crude Fat

Crude fat was determined by the Soxhlet extraction method (AOAC 920.39). Approximately 2 g of dried sample was extracted continuously with petroleum ether (boiling point 40-60°C) for 6-8 hours. The solvent was evaporated, and the extracted lipid was dried and weighed. Fat content was expressed as a percentage of dry weight. Ash Content: Ash content was determined by incinerating approximately 2 g of dried sample in a muffle furnace at 550°C for 6 hours until a light grey or white ash was obtained. The residue was cooled in a desiccator and weighed.

##### Crude Fibre

Crude fibre was determined following AOAC (2019). Defatted samples were successively digested with dilute sulfuric acid and sodium hydroxide under controlled heating. The residue was filtered, dried, weighed, incinerated, and reweighed. Crude fibre content was calculated from the loss in weight after ashing.

##### Total Carbohydrate

Total carbohydrate was determined using the phenol-sulfuric acid colorimetric method. Approximately 100 mg of dried mushroom powder was hydrolysed, reacted with 5% phenol and concentrated sulfuric acid, and incubated for colour development. Absorbance was measured at 490 nm using a UV-Visible spectrophotometer. Glucose was used for preparation of the calibration curve, and carbohydrate concentration was expressed as g/100 g dry weight.

##### Energy Value

The metabolizable energy value of each sample was calculated using the Atwater conversion factors:

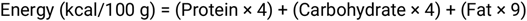

where protein, carbohydrate, and fat contents are expressed as g/100 g dry weight.

#### 2.4.2 Mineral Analysis

Mineral composition was determined according to AOAC (2019) procedures.

Approximately 0.5 g of dried mushroom powder was digested with a nitric acid-perchloric acid (HNO_3_,HClO_4_) mixture until a clear solution was obtained. The digest was cooled, filtered through Whatman No. 1 filter paper, and diluted to 50 mL with deionized water. The concentrations of calcium (Ca), potassium (K), sodium (Na), magnesium (Mg), iron (Fe), zinc (Zn), copper (Cu), manganese (Mn), and selenium (Se) were determined using Atomic Absorption Spectrophotometry (AAS) following calibration with certified standard solutions. Phosphorus (P) was determined spectrophotometrically using the molybdenum blue method at 660-880 nm after appropriate colour development (AOAC, 2019).

Mineral concentrations were expressed as mg/100 g dry weight.

#### 2.4.3 Vitamin Analysis

Vitamin analysis was performed using standard analytical procedures.

##### Determination of B-Complex Vitamins and Vitamin D2

Approximately 1 g of dried mushroom powder was extracted using appropriate extraction solvents under light-protected conditions. The extracts were filtered through a 0.45-μm membrane filter before injection into a High-Performance Liquid Chromatography (HPLC) system equipped with a reverse-phase C18 column and UV detector. Identification and quantification of vitamins were achieved by comparing the retention times and peak areas of samples with those of certified vitamin standards. Calibration curves were prepared for each vitamin, and concentrations were calculated accordingly.

##### Determination of Vitamin C

Vitamin C (ascorbic acid) was determined using the 2,6-dichlorophenolindophenol (DCPIP) titrimetric method. Approximately 10 g of mushroom sample was homogenized in 3% metaphosphoric acid, filtered, and titrated against standardized DCPIP dye until a light pink endpoint persisted for 15 seconds. Vitamin C content was calculated from the titre value using the standardization factor and expressed as mg/100 g (AOAC, 2019).

#### 2.4.4 Quality Assurance and Quality Control

All glassware was acid-washed and rinsed thoroughly with deionized water before use. Analytical-grade reagents were used throughout the study. Standard reference materials and reagent blanks were included during mineral and vitamin analyses for quality assurance. Instrument calibration was performed before each analytical run using certified standards. All analyses were conducted in triplicate, and the results were reported as mean ± standard deviation (SD). Statistical analyses were performed using SPSS, with significant differences among treatment means determined by one-way ANOVA followed by Tukey’s post hoc test at p < 0.05.

### 2.5 Sample Extraction and Mycochemical Profiling

#### 2.5.1 Preparation of Extracts

Fruiting bodies were oven-dried at 60°C to constant weight, milled to fine powder (0.5 mm sieve) and stored in airtight containers at 4°C until analysis. Dried powder (20 g) was extracted separately with 200 mL of 70% ethanol (v/v) and distilled water (1:10 w/v) in 250 mL Erlenmeyer flasks. Extraction was performed on an orbital shaker at 150 rpm, 25 ± 2°C for 24 h. The mixture was filtered through Whatman No. 1 filter paper and the filtrate concentrated under reduced pressure using a rotary evaporator at 40°C. The concentrated extracts were dried in a desiccator, weighed to determine percentage yield, and reconstituted in respective solvents to desired concentrations (Ferreira *et al*., 2009; Ferraro *et al*., 2021). Extracts were stored at -20°C until further analysis. All extractions were done in triplicate.

#### 2.5.2 HPLC Profiling of Mycochemicals

Identification and quantification of mycochemicals by HPLC were performed according to methods described for mushroom phenolics and secondary metabolites (Ferraro et al., 2021; Ferreira et al., 2009). The mycochemical constituents (phenols, flavonoids, alkaloids, saponins, tannins and terpenoids) were profiled using High Performance Liquid Chromatography (Shimadzu Nexera X2, Japan) equipped with a C18 reverse-phase column (250 × 4.6 mm, 5 μm) and UV-Vis detector. Mobile phase consisted of (A) 0.1% formic acid in water and (B) acetonitrile in gradient elution: 0-5 min 10% B, 5-20 min 10-50% B, 20-30 min 50-90% B, 30-35 min 90-10% B. Flow rate was 1.0 mL/min, injection volume 20 μL, column temperature 30°C and detection at 280 nm. Compounds were identified by comparing retention times and UV spectra with authentic standards and quantified using calibration curves. Results were expressed as mg/g dry weight.

#### 2.5.3 Determination of Total Phenolic Content (TPC)

TPC was determined by Folin-Ciocalteu method. Briefly, 0.5 mL of extract (1 mg/mL) was mixed with 2.5 mL of 10% Folin-Ciocalteu reagent and incubated for 5 min. Then 2 mL of 7.5% sodium carbonate was added and incubated at 45°C for 40 min. Absorbance was read at 765 nm against blank (Singleton *et al.,* 1999). Gallic acid (20-100 μg/mL) was used as standard. TPC was expressed as mg gallic acid equivalent per gram of extract (mg GAE/g).

#### 2.5.4 DPPH Radical Scavenging Assay

The antioxidant activity was evaluated using the DPPH radical scavenging method (Brand-Williams et al., 1995). Antioxidant activity was evaluated by 1,1-diphenyl-2-picrylhydrazyl (DPPH) assay. Different concentrations of extracts (0.5-5 mg/mL) were prepared. 1 mL of each concentration was mixed with 1 mL of 0.1 mM DPPH in methanol, vortexed and incubated in darkness for 30 min at 25°C. Absorbance was measured at 517 nm. Ascorbic acid was used as positive control. Percentage radical scavenging activity (% RSA) was calculated as:

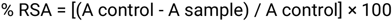

IC50, the concentration required to scavenge 50% of DPPH radicals, was determined from the dose-response curve by linear regression.

#### 2.5.5 Ferric Reducing Antioxidant Power (FRAP) Assay

FRAP assay was performed according to Benzie and Strain (1996) method. FRAP reagent was prepared by mixing 300 mM acetate buffer (pH 3.6), 10 mM TPTZ in 40 mM HCl and 20 mM FeCl3 in ratio 10:1:1. 1.5 mL of FRAP reagent was mixed with 0.1 mL of extract (1 mg/mL) and 0.1 mL distilled water, incubated at 37°C for 10 min and absorbance read at 593 nm. FeSO4 (100-1000 μmol/L) was used for calibration. Results were expressed as mmol Fe2+ equivalent per gram of extract (mmol Fe2+ eq/g).

### 2.6 Antimicrobial Activity

#### 2.6.1 Test Organisms and Inoculum Preparation

##### Six clinical pathogens

*Escherichia coi, Staphylococcus aureus, Klebsiella pneumoniae, Pseudomonas aeruginosa, Salmonella Typhi* and *Candida albicans* were obtained from State Specialist Hospital, Okitipupa (with ethical approval no: SP/OK/2026/00023). Isolates were confirmed by standard microbiological methods and maintained on Nutrient Agar at 4°C. Inoculum was standardized to 0.5 McFarland (1.5 × 10^8 CFU/mL) using sterile normal saline.

#### 2.6.2 Agar Well Diffusion Assay

Antimicrobial susceptibility testing followed standard laboratory procedures (Clinical and Laboratory Standards Institute, 2018; European Committee on Antimicrobial Susceptibility Testing, 2023). Antimicrobial activity was evaluated by agar well diffusion method on Mueller-Hinton Agar (for bacteria) and Sabouraud Dextrose Agar (for Candida). 100 μL of standardized inoculum was spread on agar plates. Wells of 6 mm diameter were bored aseptically and filled with 100 μL of extract at 5 mg/mL. Ciprofloxacin (5 μg/disc) for bacteria and fluconazole (25 μg/disc) for fungi were used as positive controls, while solvents (70% ethanol and distilled water) served as negative controls. Plates were incubated at 37°C for 24 h (bacteria) and 28°C for 48 h (fungus). Zones of inhibition were measured in mm using a transparent ruler. Assays were in triplicate.

#### 2.6.3 Minimum Inhibitory Concentration (MIC)

The minimum inhibitory concentration was done followed standard laboratory procedures (CLSI, 2018; EUCAST, 2023). MIC was determined by broth microdilution method in 96-well microtiter plates. Two-fold serial dilutions of extracts (100 to 6.25 mg/mL) were prepared in Mueller-Hinton Broth. 100 μL of standardized inoculum was added to each well. Plates were incubated at 37°C for 24 h. The lowest concentration with no visible growth was taken as MIC.

### 2.6 Data analysis

All analyses were in triplicate. Data expressed as mean ± SD. One-way ANOVA with Tukey’s HSD test at p<0.05 was used for cultivation, nutritional and bioactivity data using SPSS v25 and GraphPad Prism v9.

## 3.0 Results

### 3.1 Cultivation Performance

Growth performance and biological efficiency. Cassava peels exhibited the shortest spawn run (10.1 ± 0.3 d), pinhead formation (3.5 ± 0.1 d) and fruiting time (13.2 ± 0.3 d), while Gmelina sawdust was slowest (12.2 ± 0.5, 6.9 ± 0.2, 17.0 ± 0.3 d, respectively) (Table 1).Conversely, biological efficiency (BE) was highest on oil palm pressed fibre (74.2 ± 2.0%), followed by Gmelina sawdust (68.1 ± 2.5%) and cassava peels (59.3 ± 2.6%) (Table 2). These underscored their value as low-cost, sustainable cultivation material, while simultaneously addressing oil palm fruit fibre and cassava wastes disposal problems in Nigeria.

**Table 1:**
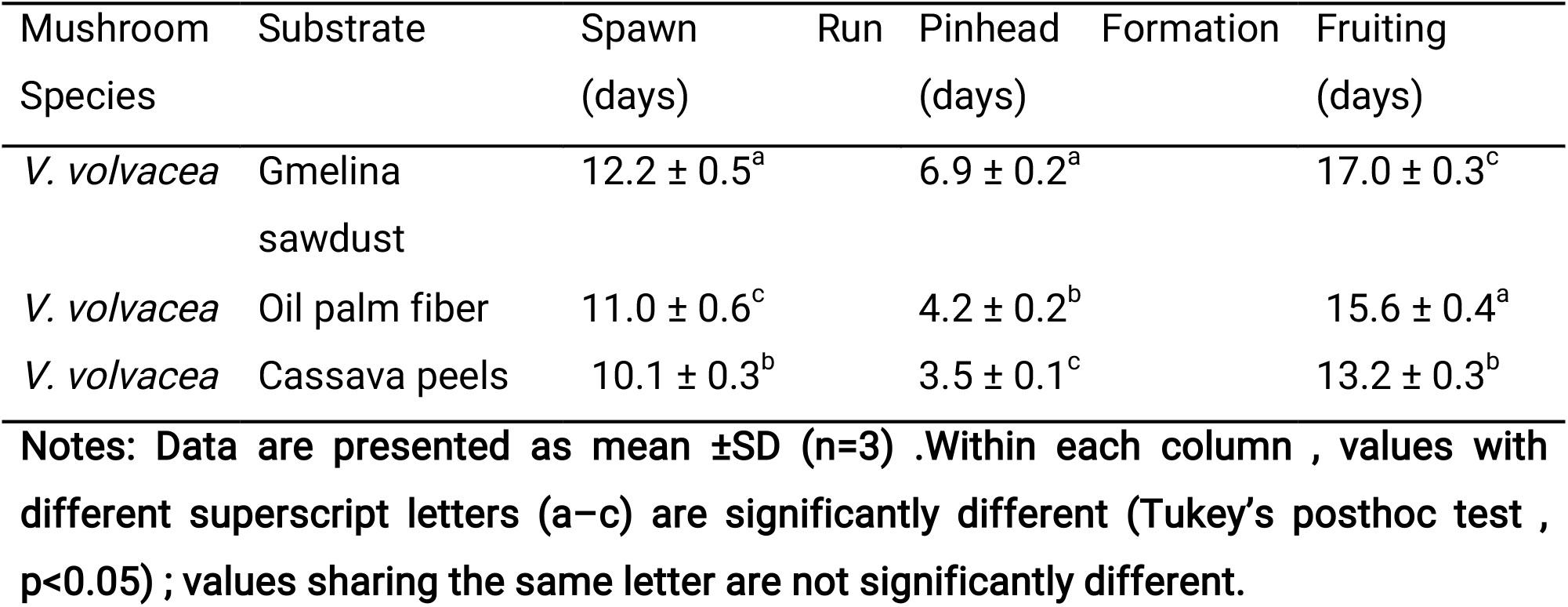
Spawn Run and Fruiting Time of Mushrooms on Different Substrates.

**Table 2:** Biological Efficiency (%) of *V. volvaceae* on Different Substrates.

| Species | Substrate | BE (%) ± SD |
| --- | --- | --- |
| *V. volvacea* | Gmelina sawdust | 68.1 ± 2.5<sup>a</sup> |
| *V. volvacea* | Oil palm pressed fiber | 74.2 ± 2.0<sup>b</sup> |
| *V. volvacea* | Cassava peels | 59.3 ± 2.6<sup>c</sup> |

Notes: Data are presented as mean ± SD (n = 3). Within each column, values with different superscript letters (a–c) are significantly different (Tukey’s post hoc test, p < 0.05); values sharing the same letter are not significantly different.

### 3.2 Molecular Identification

Morphological and microscopic analyses provided preliminary identification as shown in Table 3. Molecular identification, Internal transcribed spacer (ITS) sequence showed 100% query coverage and 98.5% identity with *V. volvacea* MG280840.1, with LSU 98.7% and RPB2 98.9% (Table 4), confirming taxonomic placement. ITS is the universal fungal barcode and GenBank BLASTn ≥98% identity is accepted for species delimitation. The 1.1-1.5% divergence likely represents intraspecific variation in African isolates. This multi-locus approach provides robust identification beyond morphology and contributes African *V. volvacea* sequences to global databases. The gene sequence was deposited in the GenBank with accession number PX630179.1.

**Table 3:**
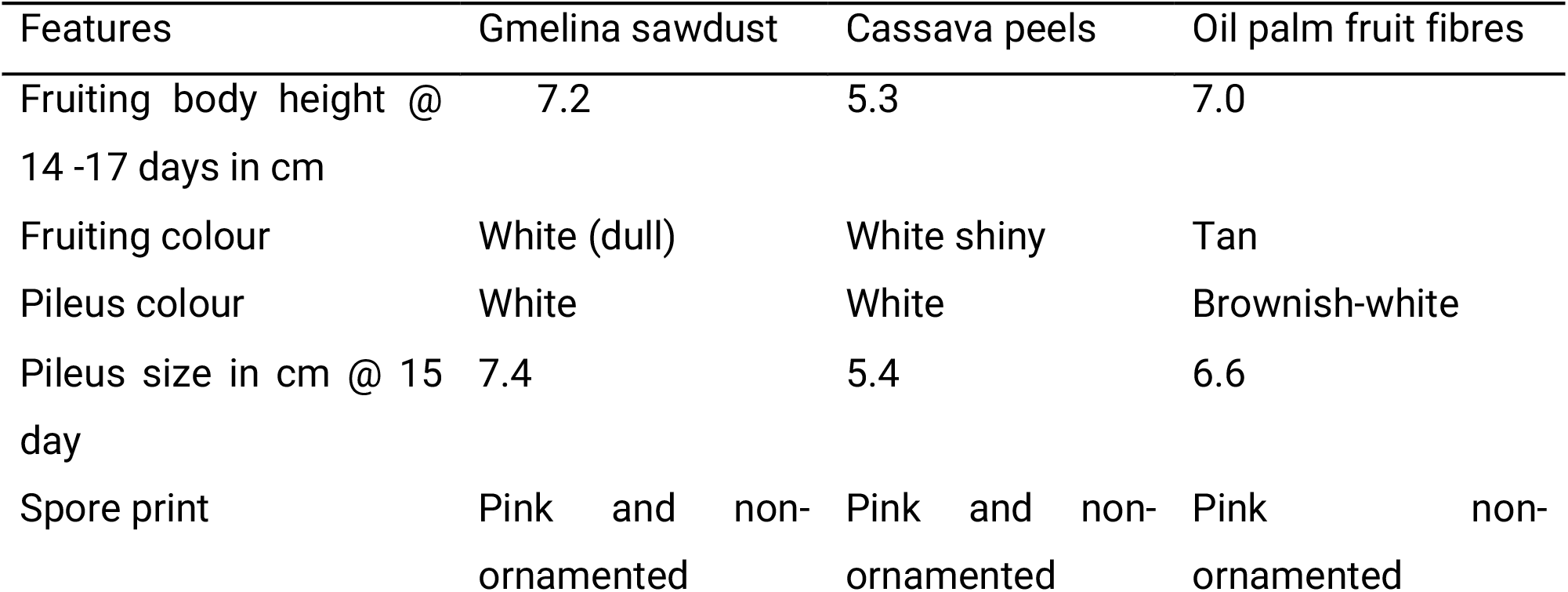
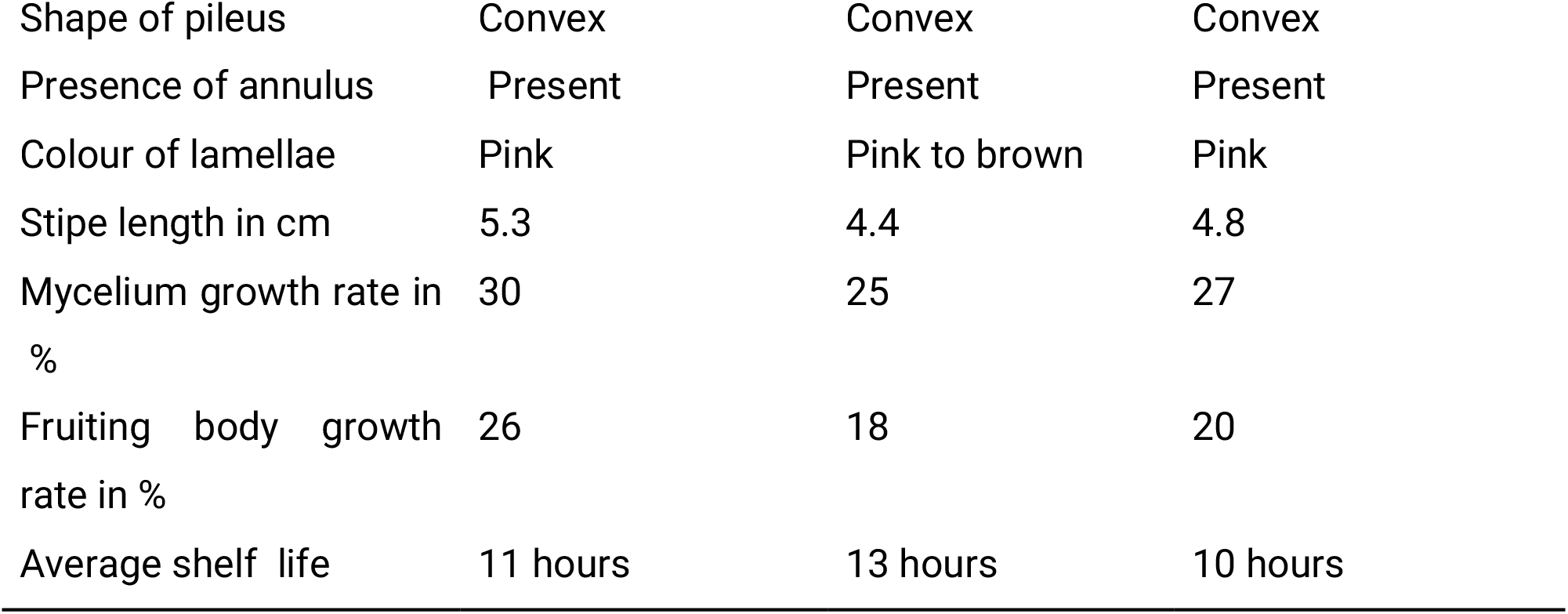
Features of the fruiting bodies of *Volvariella volvacae* cultivated on Gmelina sawdust, Cassava peels and Oil palm fruit pressed fibres respectively.

| Features | Gmelina sawdust | Cassava peels | Oil palm fruit fibres |
| --- | --- | --- | --- |
| Fruiting body height @ 14 -17 days in cm | 7.2 | 5.3 | 7.0 |
| Fruiting colour | White (dull) | White shiny | Tan |
| Pileus colour | White | White | Brownish-white |
| Pileus size in cm @ 15 day | 7.4 | 5.4 | 6.6 |
| Spore print | Pink and non-ornamented | Pink and non-ornamented | Pink non-ornamented |
| Shape of pileus | Convex | Convex | Convex |
| Presence of annulus | Present | Present | Present |
| Colour of lamellae | Pink | Pink to brown | Pink |
| Stipe length in cm | 5.3 | 4.4 | 4.8 |
| Mycelium growth rate in % | 30 | 25 | 27 |
| Fruiting body growth rate in % | 26 | 18 | 20 |
| Average shelf life | 11 hours | 13 hours | 10 hours |

**Table 4:** Molecular identification summary of *Volvariella volvacae* (BLASTn)

| Species | Locus | Identity (%) | Query Coverage (%) |
| --- | --- | --- | --- |
| *V. volvacea* | ITS | 98.5 | 100 |
| *V. volvacea* | LSU | 98.7 | 99 |
| *V. volvacea* | RPB2 | 98.9 | 97 |

**Abbreviations:** ITS: Internal transcribed spacer region; LSU: Large subunit ribosomal RNA gene; RPB2: RNA polymerase II second-largest subunit gene.

### 3.4 Nutritional Composition

Fruiting bodies were rich in protein (10.2-12.3 g/100g dry weight, Figure 1), low in fat, and high in fibre, ash and carbohydrate, consistent with the established low-fat, high-protein profile of edible mushrooms. Mineral analysis (Figure 2) revealed phosphorus as the dominant mineral (>20%), followed by K, Ca, Fe and Zn, this mineral density, particularly Fe and Zn, is nutritionally significant for combating micronutrient deficiencies in developing countries. Vitamin B-complex and D2 were detected (Figure 3), confirming mushrooms as one of the few non-animal sources of vitamin D2. The significant substrate effect - Gmelina sawdust enriching B-complex while oil palm fibre enriched D2 - indicates substrate-dependent vitamin biosynthesis, possibly linked to lignin degradation precursors. D2.

**Figure 1:**
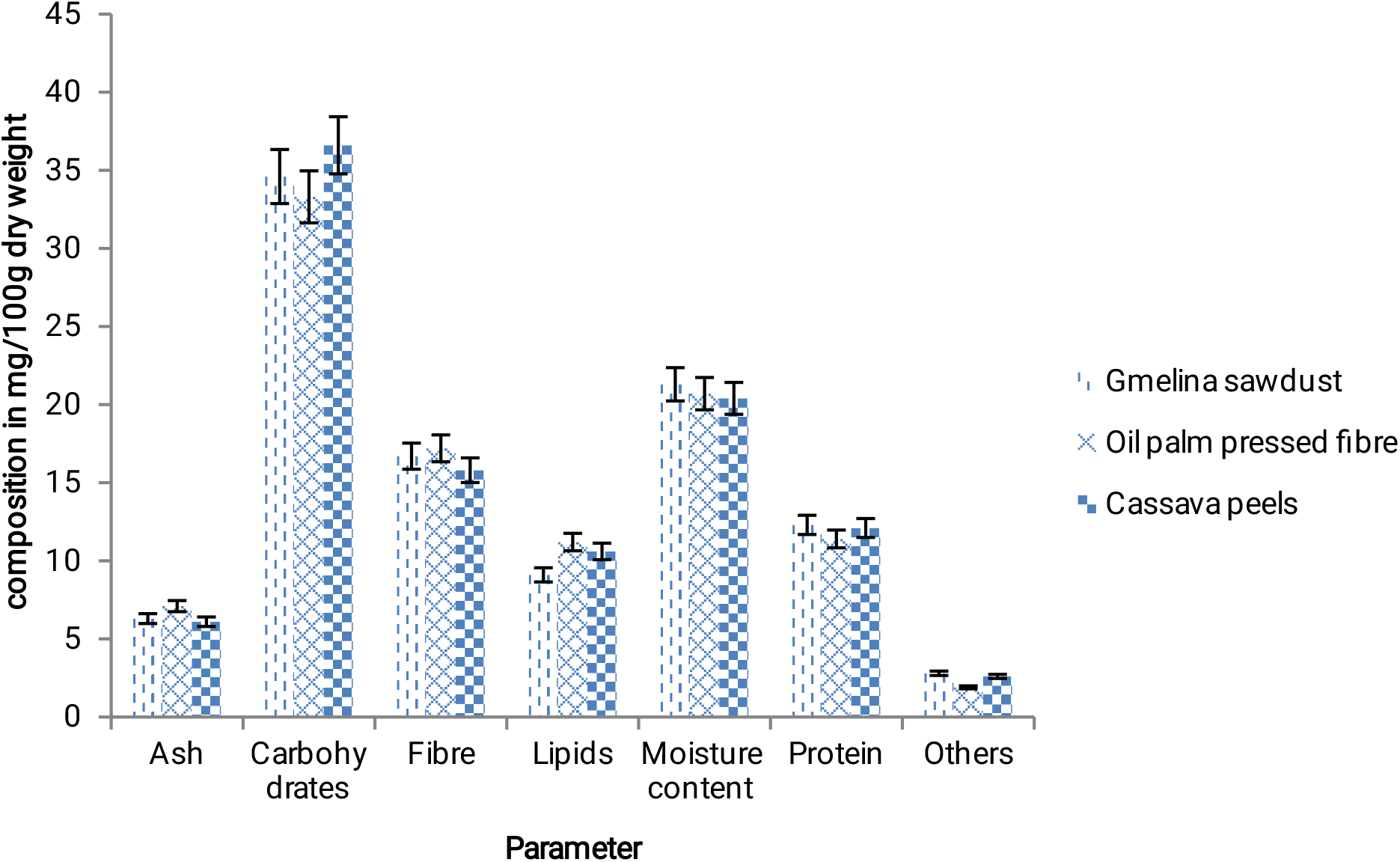
Proximate composition of V. *volvacea* grown on different agro-wastes (mg/100g dry weight)

**Figure 2:**
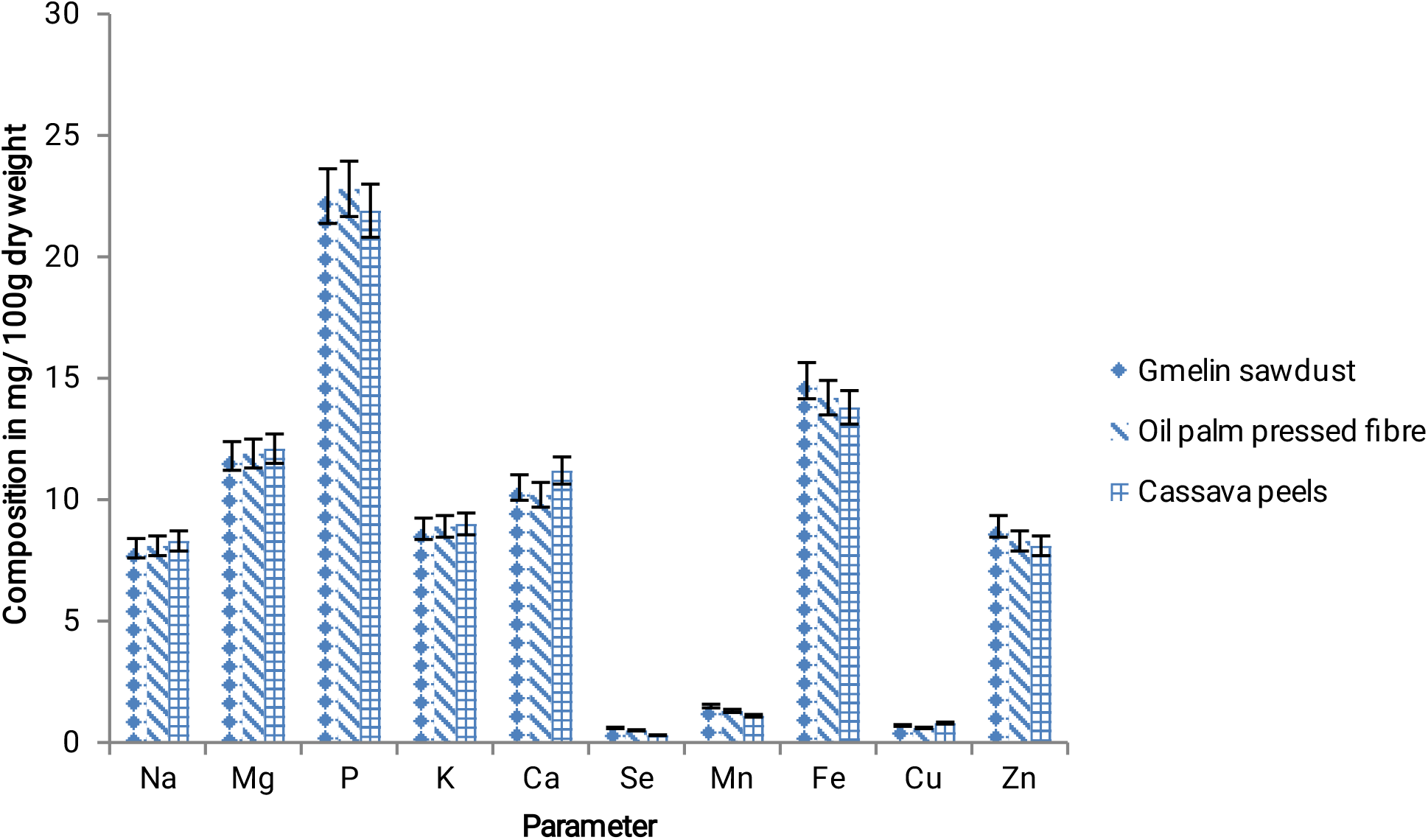
Minerals composition of *V. volvacea* grown on different agro-wastes mg/100g dry weight

**Figure 3:**
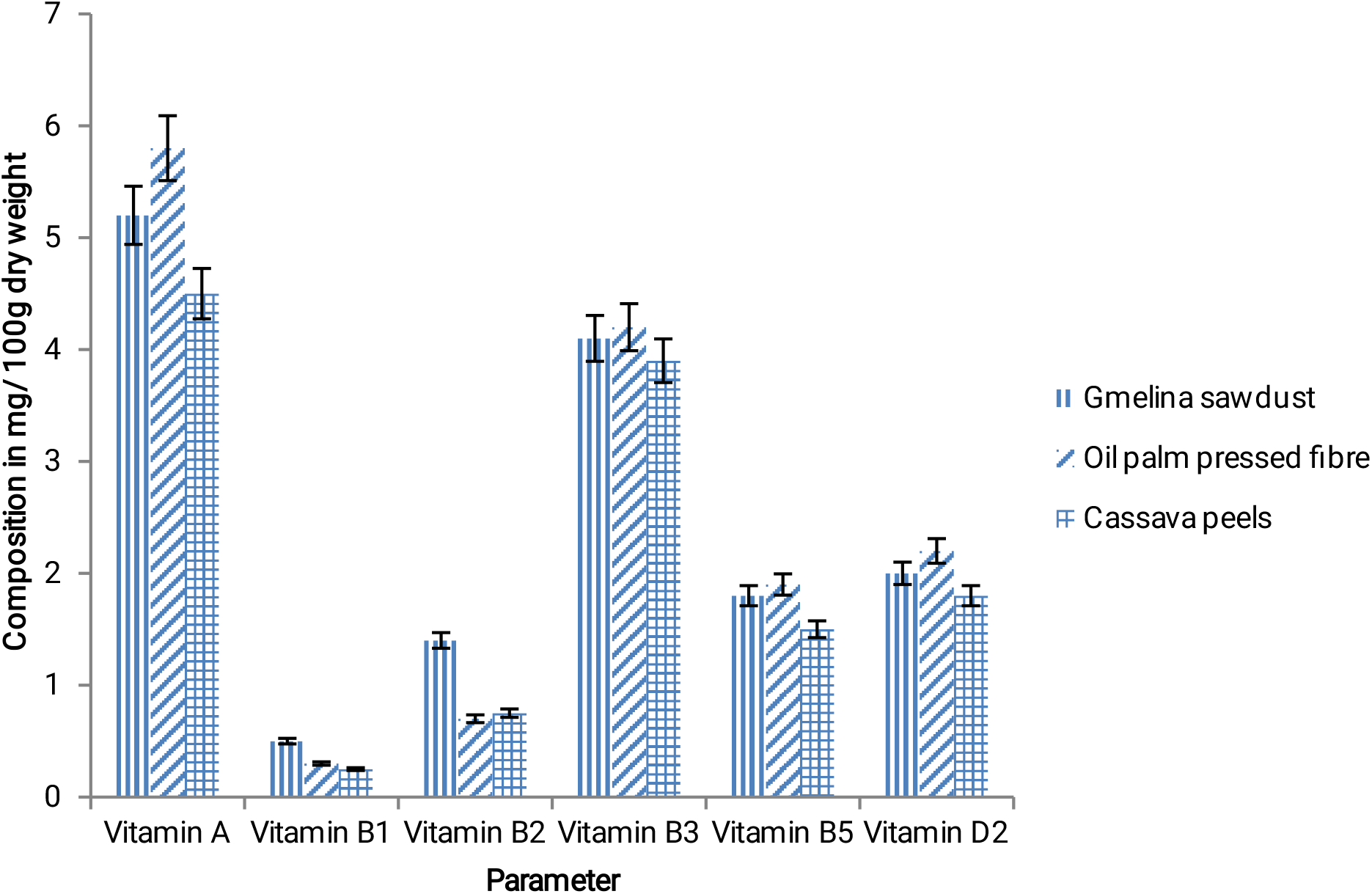
Vitamins composition of *V. volvacea* grown on different agro-wastes mg/ 100g dry weight

### 3.5 Substrate-Dependent Variation in Mycochemical Composition and Antioxidant Activity

Substantial variation in mycochemicals was observed (Figure 4). Alkaloids (14.3 mg/100g) and tannins (12.6 mg/100g) dominated, with phenols, saponins-like compounds, flavonoids and terpenoids present. This profile indicates that cultivation substrate modulates mycochemical biosynthesis. Cassava peel-grown mushrooms (VVCP) had highest phenolics, saponins-like compounds, alkaloids and tannins, obtained from secondary metabolism from residual cyanogenic glycosides. This translated to enhanced antioxidant activity. At 5 mg/mL, DPPH RSA ranged 47.40% (VVGS, IC50 4.65 mg/mL) to 53.21% (VVCP, IC50 3.80 mg/mL) (Table 5), FRAP 10.10-10.20 mmol Fe2+ eq/g (Table 6) and TPC 1.50-1.65 mg GAE/g (Table 7), with VVCP highest. The strong positive correlation between TPC and FRAP (r=0.99) and negative correlation between TPC/FRAP and DPPH IC50 (Figure 5) confirms that phenolic compounds are major contributors to antioxidant capacity. Although our IC50 values indicate moderate antioxidant activity compared to ascorbic acid, typical for crude extracts.

**Figure 4:**
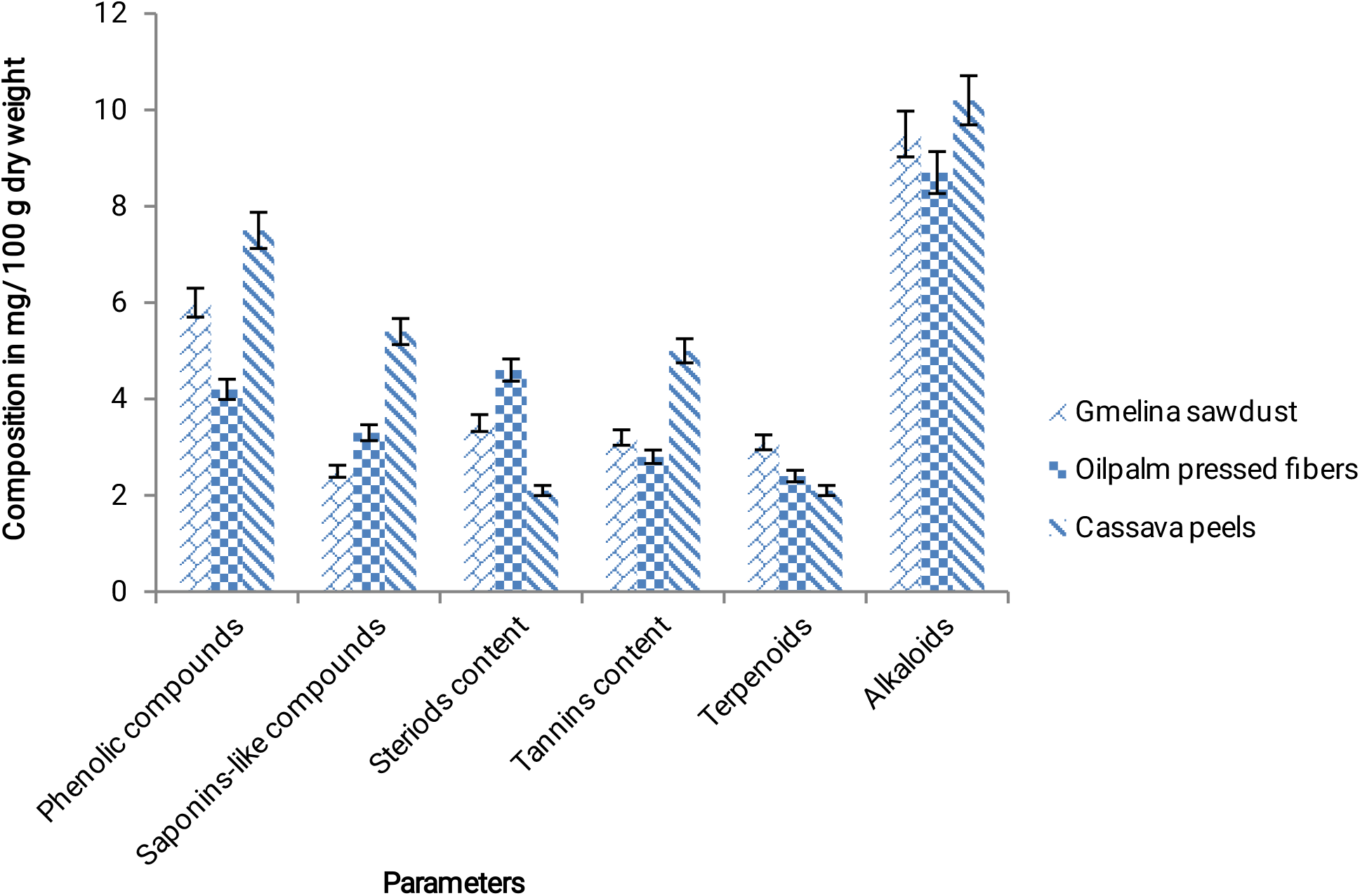
Mycochemicals content in *V. volvacea* grown on different agro wastes in mg/ 100g dry weight

**Table 5:** Antioxidant (DPPH assay) values of *V. volvacea* (5mg/ml)

| Species | Substrate | RSA % | IC<sub>50</sub> (5mg/ml) |
| --- | --- | --- | --- |
| *Volvariella volvacea* | Sawdust | 47.40± 0.3 | 4.65± 0.4 |
| *Volvariella volvacea* | Cassava Peels | 53.21± 0.7 | 3.80± 0.5 |
| *Volvariella volvacea* | Oil palm Fiber | 50.34± 0.4 | 4.00± 0.2 |

**Note:** The inverse relationship between RSA and IC<sub>50</sub> values further supports the greater antioxidant capacity of the extracts derived from the cassava peel substrate.
**Abbreviations:** IC<sub>50</sub>: Half maximal inhibitory concentration; RSA: Radical-scavenging activity

**Table 6:** Antioxidant (FRAP assay) values of *V. volvacea* (5mg/ml)

| Species | Substrate | AAE (mmol/g) | Trolox (mmol TE/g) |
| --- | --- | --- | --- |
| *Volvariella volvacea* | Sawdust | 10.10± 0.4<sup>a</sup> | 9.45±0.1<sup>b</sup> |
| *Volvariella volvacea* | Cassava Peels | 10.20±0.2<sup>c</sup> | 9.50±0.3<sup>a</sup> |
| *Volvariella volvacea* | Palm Fiber | 10.17±0.1<sup>b</sup> | 9.48±0.2<sup>c</sup> |

**Note:** Both the ascorbic acid calibration curve and Trolox calibration curve were used to determine FRAP values. Abbreviations: AAE: Ascorbic acid equivalent; FRAP: Ferric reducing antioxidant power; TE: Trolox equivalents. FRAP values showed slight but statistically significant differences among substrates ( $p < 0.05$ ). Mushrooms cultivated on cassava peels recorded the highest antioxidant reducing power (10.20 mmol/g AAE and 9.50 mmol TE/g), followed by oil palm fiber and Gmelina sawdust. Although numerical differences were relatively small, ANOVA indicated significant variation among substrates. Notes: Data are presented as mean ± SD ( $n = 3$ ), within each column, values with different superscript letters (a–c) are significantly different

**Table 7:** Total Phenolic Coentent (TPC) values of *V. volvacea* (5mg/ml)

| Species | Substrate | mg GAE/g | mg CAE |
| --- | --- | --- | --- |
| *Volvariella volvacea* | Sawdust | 1.62± 0.3 | 10.75± 0.6 |
| *Volvariella volvacea* | Cassava Peels | 1.65± 0.5 | 10.95± 0.2 |
| *Volvariella volvacea* | Palm Fiber | 1.50± 0.3 | 10.62± 0.4 |

**Abbreviations:** CAE: Catechin equivalents; GAE: Gallic acid equivalents.
**Notes:** Data are presented as mean ± SD ( $n = 3$ ). Total phenolic content varied significantly with cultivation substrate ( $p < 0.05$ ). Cassava peels yielded the highest phenolic concentration (1.65 mg GAE/g and 10.95 mg CAE), followed by Gmelina sawdust (1.62 mg GAE/g), while oil palm fiber recorded the lowest phenolic content (1.50 mg GAE/g). The increased phenolic content in mushrooms cultivated on cassava peels corresponded with their superior antioxidant activity.

**Figure 5:**
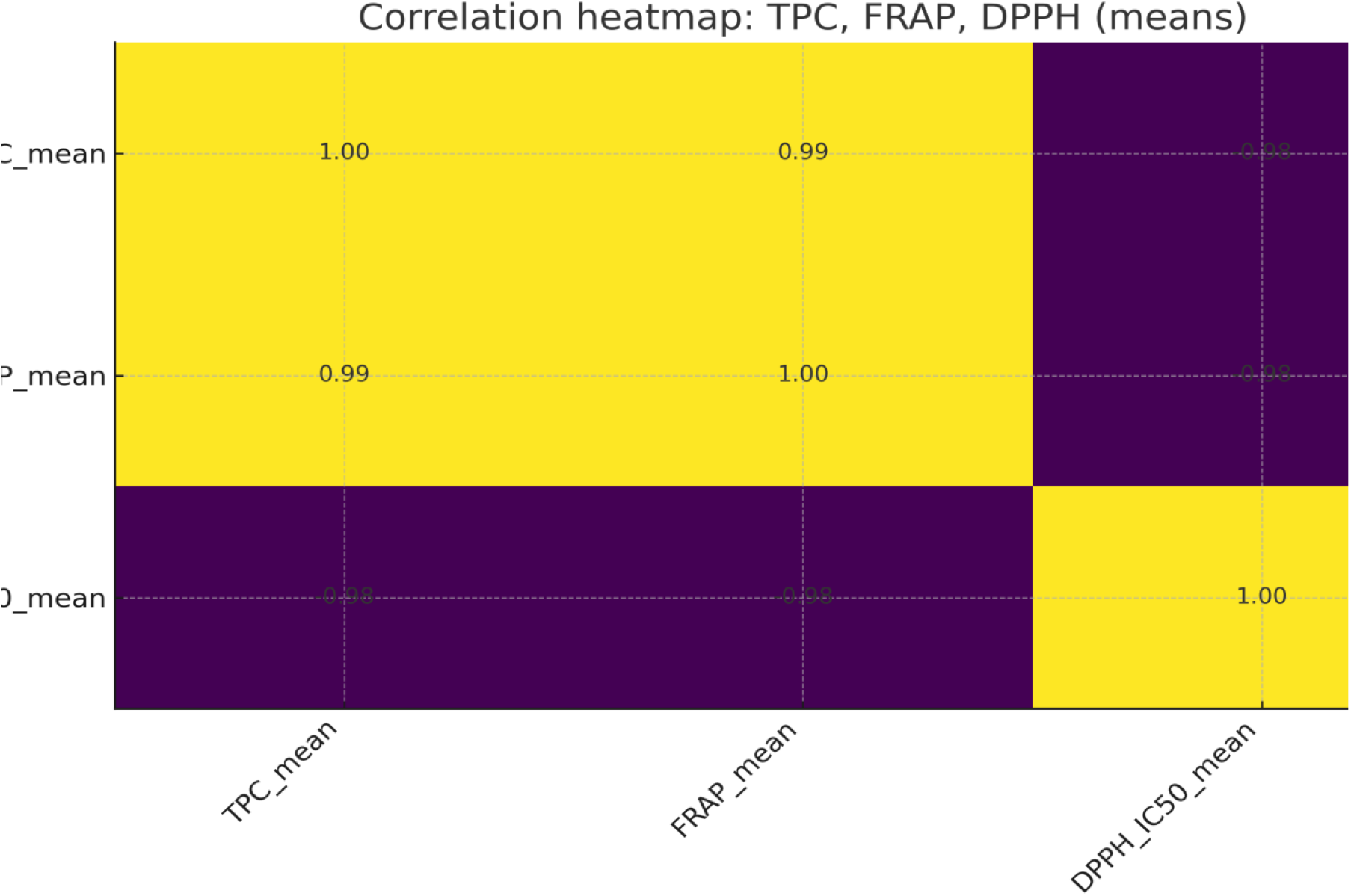
Pearson correlation heatmap of mean Total Phenolic Content (TPC), Ferric Reducing Antioxidant Power (FRAP) and DPPH radical scavenging IC50. Yellow indicates strong positive correlation (r ∼ +1.0) and dark purple indicates strong negative correlation (r ∼ -1.0).

### 3.7 Antimicrobial Properties

Ethanolic extracts showed broader activity (zones 15.1-17.4 mm) than aqueous extracts (11.9-13.3 mm) against six clinical pathogens (Table 8), consistent with higher solubility of phenolics and alkaloids in ethanol. Highest activity against S. aureus (17.4 ± 0.6 mm) and E. coli (16.7 ± 0.5 mm) with MIC of 25 mg/mL for E. coli and C. albicans and 50 mg/mL for others (Table 9) demonstrates broad-spectrum potential, though lower than ciprofloxacin/fluconazole controls per CLSI and EUCAST breakpoints. The results validate the traditional ethnomedicinal use of V.volvacea and highlight its potential as natural antimicrobial agents.

**Table 8:** Antimicrobial Activity of *V. volvacea* extracts (Zone of Inhibition in mm at 5 mg/ml)

| Pathogen | *V. volvacea* (ET) | *V. volvacea* (Aq) | Positive Control | Negative control |
| --- | --- | --- | --- | --- |
| *E. coli* | 16.7 ± 0.5<sup>a</sup> | 12.8 ± 0.3<sup>c</sup> | 25.1 ± 0.4<sup>b</sup> | 5.0 ± 0.4<sup>d</sup> |
| *S. aureus* | 17.4 ± 0.6<sup>b</sup> | 13.3 ± 0.4<sup>a</sup> | 27.5 ± 0.5<sup>c</sup> | 6.2 ± 0.1<sup>d</sup> |
| *K. pneumonia* | 15.9 ± 0.4<sup>d</sup> | 12.2 ± 0.3<sup>c</sup> | 24.6 ± 0.4<sup>b</sup> | 6.0 ± 0.2<sup>a</sup> |
| *P. aeruginosa* | 15.1 ± 0.4<sup>a</sup> | 11.9 ± 0.3<sup>c</sup> | 23.8 ± 0.4<sup>b</sup> | 5.5 ± 0.3<sup>d</sup> |
| *S. typhi* | 16.5 ± 0.5<sup>a</sup> | 12.7 ± 0.3<sup>c</sup> | 25.4 ± 0.5<sup>b</sup> | 6.1 ± 0.1<sup>d</sup> |
| *C. albicans* | 15.8 ± 0.4<sup>c</sup> | 12.4 ± 0.3<sup>d</sup> | 26.3 ± 0.4<sup>a</sup> | 5.2 ± 0.2<sup>b</sup> |

ET =ethanol Aq = aqueous
Notes: Data are presented as mean ± SD (n = 3). Within each row, values with different superscript letters (a–d) are significantly different (Tukey's post hoc test, p < 0.05); values sharing the same letter are not significantly different. The negative control was 20.0% (v/v) ethanol. Ciprofloxacin and fluconazole were used as positive controls for bacteria and fungi, respectively

**Table 9:**
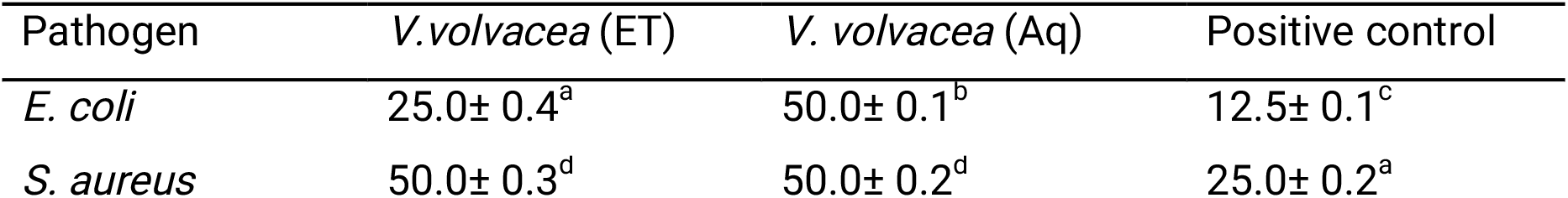
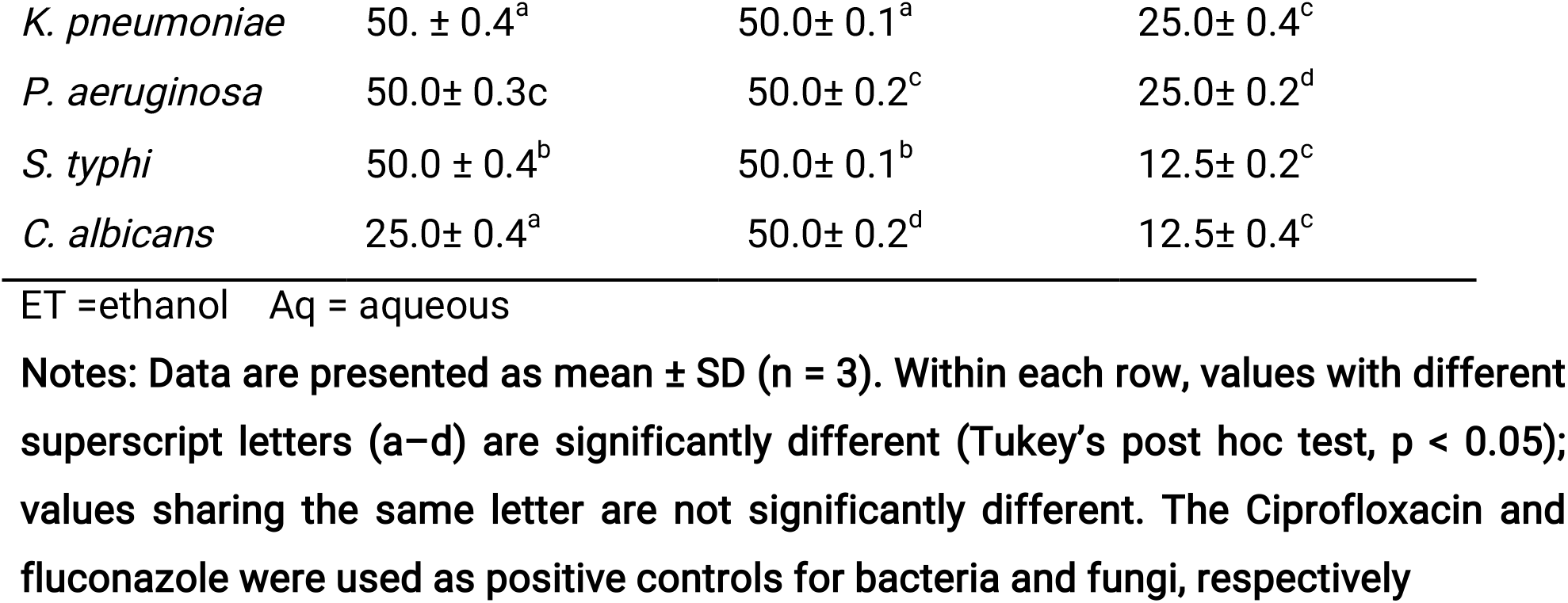
MIC of *V. volvacea* extract against clinical pathogens (mg/ml)

| Pathogen | *V. volvacea* (ET) | *V. volvacea* (Aq) | Positive control |
| --- | --- | --- | --- |
| *E. coli* | 25.0 ± 0.4<sup>a</sup> | 50.0 ± 0.1<sup>b</sup> | 12.5 ± 0.1<sup>c</sup> |
| *S. aureus* | 50.0 ± 0.3<sup>d</sup> | 50.0 ± 0.2<sup>d</sup> | 25.0 ± 0.2<sup>a</sup> |
| *K. pneumoniae* | 50. ± 0.4<sup>a</sup> | 50.0± 0.1<sup>a</sup> | 25.0± 0.4<sup>c</sup> |
| *P. aeruginosa* | 50.0± 0.3<sup>c</sup> | 50.0± 0.2<sup>c</sup> | 25.0± 0.2<sup>d</sup> |
| *S. typhi* | 50.0 ± 0.4<sup>b</sup> | 50.0± 0.1<sup>b</sup> | 12.5± 0.2<sup>c</sup> |
| *C. albicans* | 25.0± 0.4<sup>a</sup> | 50.0± 0.2<sup>d</sup> | 12.5± 0.4<sup>c</sup> |

ET =ethanol Aq = aqueous
**Notes:** Data are presented as mean ± SD (n = 3). Within each row, values with different superscript letters (a–d) are significantly different (Tukey’s post hoc test, p < 0.05); values sharing the same letter are not significantly different. The Ciprofloxacin and fluconazole were used as positive controls for bacteria and fungi, respectively

## Discussion

The present study demonstrates that abundant tropical agricultural wastes can be effectively utilized as sustainable substrates for cultivation of *Volvariella volvacea,* while simultaneously influencing mushroom productivity, nutritional quality and bioactive properties (Chang and Miles 2004; Boa 2004; Stamets 2000; Albertó 2021). Although all three substrates supported successful mycelial colonization and fruiting, significant differences in agronomic performance and biochemical composition indicate that substrate characteristics play a fundamental role in regulating fungal growth and metabolism (Sánchez 2010; Silva *et al*. 2002; Ragunathan and Swaminathan 2003). These findings reinforce that cultivation substrate functions not only as physical support but also as major determinant of nutrient availability, extracellular enzyme production and secondary metabolite biosynthesis (Chen *et al*. 2016; Ferraro *et al*. 2021). Cassava peels supported fastest spawn run, whereas oil palm pressed fibre produced highest biological efficiency, demonstrating that rapid mycelial colonization does not necessarily translate into greater yield. Rapid colonization on cassava peels is likely associated with readily metabolizable carbohydrates that promote early establishment (Ragunathan and Swaminathan 2003; Fakoya *et al*. 2020; Lavelli et al. 2018). In contrast, oil palm fibre appears to provide more balanced nutritional environment capable of sustaining prolonged activity and efficient conversion into fruiting bodies (Barros *et al*. 2008; Caglarirmak 2007; Reis *et al*. 2012). Such observations emphasize that biological efficiency is governed by interacting factors including cellulose accessibility, mineral availability and gradual nutrient release rather than colonization speed alone (Silva *et al*. 2002; Sánchez 2010). Similar relationships have been reported for *V. volvacea* and other mushrooms, where lignocellulosic composition and carbon-to-nitrogen ratio substantially influence productivity (Chang and Miles 2004; Shirur *et al*. 2021; Hasan and Abdulhadi 2022). Successful cultivation on all agro-wastes highlights considerable potential of residue valorization within tropical systems. Nigeria generates substantial quantities of cassava peels, sawdust and oil palm residues annually, much of which remains underutilized (Adedokun *et al*. 2022; Boa 2004). Their conversion into edible mushrooms provides environmentally friendly strategy for reducing agricultural waste while producing nutritious food and value-added bioactive products. This circular bioeconomy approach aligns with global efforts to improve food security and promote sustainable production through efficient utilization of renewable biomass (Lavelli *et al*. 2018; Andrade *et al*. 2024; Mayirnao *et al*. 2024).

Accurate species identification remains fundamental for mushroom biotechnology because morphological characteristics alone are often insufficient (Cai and Hyde 2019; Schoch *et al*. 2012; Adetuwo *et al*., 2026). In the present study, molecular analyses confirmed identity as V. volvacea with high sequence similarity using ITS, LSU and RPB2 regions. Combined use of multiple markers provides greater taxonomic confidence than single barcode region and minimizes misidentification (White *et al*. 1990; Benson et al. 2013; Schoch *et al*. 2012). Multi-locus identification has become increasingly important because it improves phylogenetic resolution (Cai and Hyde 2019; Hasan and Abdulhadi 2022). Furthermore, deposition of sequence data in GenBank contributes valuable information for African mushroom germplasm where genomic resources remain limited (Benson *et al*. 2013).

Nutritional evaluation further demonstrated that *V. volvacea* constitutes highly valuable functional food. Regardless of substrate, mushrooms contained appreciable protein, carbohydrates, fibre and ash while maintaining low lipid content, consistent with well-established nutritional profile of edible mushrooms (Aremu *et al*. 2018; Barros et al. 2008; Reis *et al*. 2012; Roupas *et al*. 2012; Valverde *et al*. 2015). Higher protein on Gmelina sawdust suggests substrate composition can influence nitrogen assimilation, while mushrooms on cassava peels accumulated greater carbohydrates, reflecting differences in carbon availability (Ragunathan and Swaminathan 2003; Aremu et al. 2018; AlAli *et al*. 2021). Consistently low fat represents nutritional advantage in context of obesity and cardiovascular disease (Rathore et al. 2017; Roupas *et al*. 2012; AlAli et al. 2021; Anachad *et al*. 2023).

Mineral analysis confirmed nutritional importance. Phosphorus was predominant, followed by magnesium, iron and calcium. These micronutrients perform indispensable physiological functions including skeletal development, energy metabolism and immune regulation (Aremu *et al*. 2018; Caglarirmak 2007; Barros *et al*. 2008; Ahmed *et al*. 2020). Favourable potassium-to-sodium ratio is desirable because diets rich in potassium and low in sodium are associated with reduced hypertension risk (Barros *et al*. 2008; Valverde *et al*. 2015; Allam 2024).

Vitamin profile further illustrates substrate influence. Detectable vitamins A, B-complex and D2 confirmed V. volvacea as important dietary source of essential micronutrients (Reis *et al*. 2012; Valverde *et al*. 2015; Rathore *et al*. 2017). Higher vitamin D2 on oil palm fibre suggests substrate composition may influence ergosterol metabolism, whereas Gmelina favoured B-complex accumulation (Sánchez 2010; Ferreira *et al*. 2009). Such targeted nutritional enhancement presents opportunities for functional food markets (AlAli *et al*. 2021; Ali *et al*. 2019; Andrade *et al*. 2024).

Substrate-dependent variation in mycochemical composition indicates nutritional environment influences secondary metabolism (Ferraro *et al*. 2021; Guzmán 2008). Mushrooms on Gmelina accumulated higher phenolics, whereas cassava peel-grown mushrooms contained greater alkaloids, tannins and saponins. Previous studies demonstrated that lignocellulosic substrates influence production of phenolics and terpenoids that contribute to medicinal properties (Ferraro *et al*. 2021; Ferreira *et al*. 2009; Mayirnao *et al*. 2024). Superior mycochemical profile on cassava peels was reflected in antioxidant activity. Consistency across DPPH, FRAP and TPC assays strengthens confidence in biological significance and demonstrates that antioxidant capacity is closely associated with substrate-induced changes (Brand-Williams *et al*. 1995; Singleton *et al*. 1999; Re *et al*. 1999; Reis *et al*. 2012; Effiong *et al*. 2024).

Correlation analysis clarified relationship between mycochemicals and antioxidant activity. Strong positive association between TPC and FRAP, together with negative relationship between TPC and DPPH IC50, confirms phenolic compounds constitute major contributors to antioxidant capacity via hydrogen donation mechanisms (Singleton et al. 1999; Brand-Williams *et al*. 1995; Ahmadi *et al*. 2022; Ferreira *et al*. 2009). These findings provide biochemical support for enhanced antioxidant performance of mushrooms cultivated on cassava peels (Effiong *et al*. 2024; Fakoya *et al*. 2020; Vamanu 2012). Antioxidant properties are particularly relevant because oxidative stress is implicated in chronic diseases including cardiovascular disorders and diabetes (Ahmadi *et al*. 2022; Ahmed *et al*. 2020; Allam 2024).

Antimicrobial evaluation demonstrated broad-spectrum activity against Gram-positive, Gram-negative bacteria and *Candida a/bicans*. Ethanolic extracts consistently exhibited greater inhibitory activity than aqueous extracts, indicating ethanol was more effective in extracting antimicrobial constituents (Bamisi *et al*. 2024; Vamanu 2012; Fakoya *et al*. 2020; Younis *et al*. 2015; Adetuwo *et al*., 2026). Relatively strong inhibition against Staphylococcus aureus, together with low MIC for Escherichia coli and Candida, demonstrates broad potential. Although lower than conventional agents per CLSI and EUCAST breakpoints (Clinical and Laboratory Standards Institute 2018; European Committee on Antimicrobial Susceptibility Testing 2023), crude extracts frequently exhibit moderate activity because bioactive metabolites are present at low concentrations and may act synergistically (Patel and Goyal 2012; Paterson 2006; Zaidman *et al*. 2005). Activity is likely attributable to combined action of phenolics, alkaloids, tannins and terpenoids that may interfere with microbial membranes and enzyme systems through complementary modes (Ferraro *et al*. 2021; Newman and Cragg 2020; Oyetayo 2012; Wasser 2010, 2011,2014).

Overall, this study revealed that Gmelina sawdust, oil palm pressed fibre and cassava peels are viable substrates for sustainable *V. volvacea* cultivation. Oil palm fibre optimizes biological efficiency while cassava peels accelerate cropping and enhance functional quality, providing evidence-based guidance for substrate selection. Tropical agricultural wastes can thus be valorized for protein-rich functional foods, supporting circular bioeconomy and food security (Adedokun *et al*. 2022; Lavelli *et al*. 2018; Shirur *et al*. 2021; Adetuwo *et al*., 2026).

## Limitations of the study

Although the present findings are encouraging, several limitations should be acknowledged. The study evaluated only three agro-waste substrates and a single *V. volvacea* isolate under controlled cultivation conditions, which may limit extrapolation of the findings to other strains or production systems. Furthermore, the antioxidant and antimicrobial evaluations were restricted to in vitro assays using crude extracts, and therefore the biological activities observed may not directly reflect physiological efficacy in vivo. Individual bioactive compounds were also not structurally identified beyond HPLC profiling, limiting mechanistic interpretation of the observed biological activities. Future investigations should therefore focus on comprehensive metabolomic characterization using LC-MS/MS and complementary analytical techniques, purification of active compounds, evaluation of synergistic interactions among metabolites, and in vivo validation of antioxidant and antimicrobial efficacy. Additional studies examining substrate physicochemical characteristics, enzyme activities and genotype-by-substrate interactions would further improve understanding of the mechanisms through which cultivation substrates regulate mushroom productivity and bioactive compound biosynthesis

## Recommendations

Beyond its implications for mushroom production, this work highlights the broader environmental significance of agro-waste valorization. The successful conversion of cassava peels, oil palm pressed fibre and Gmelina sawdust into nutritionally valuable mushrooms represents an effective strategy for reducing agricultural waste accumulation while generating value-added products. This approach supports the principles of sustainable agriculture and the circular bioeconomy by transforming low-value lignocellulosic residues into functional foods with demonstrated nutritional and bioactive benefits. For many developing countries where these agricultural residues are abundant, mushroom cultivation offers an economically feasible strategy for improving food security, generating rural income and reducing environmental pollution associated with open-field disposal or burning of agricultural wastes.

## Conclusion

This study provides compelling evidence that tropical agricultural wastes constitute valuable resources for the sustainable cultivation of *Volvariella volvacea.* Beyond supporting efficient mushroom production, these substrates significantly influence nutritional composition, micronutrient accumulation and the biosynthesis of biologically active compounds. The integration of molecular authentication, nutritional evaluation, antioxidant assessment and antimicrobial analysis provides a comprehensive understanding of the functional value of *V. volvacea* cultivated on locally available agro-wastes. These findings strengthen the scientific basis for promoting agricultural waste valorization through mushroom cultivation and support the development of environmentally sustainable production systems capable of supplying nutrient-rich functional foods and promising sources of natural bioactive compounds for future food and nutraceutical applications.

## Acknowledgements

The author welcomes collaboration from interested researchers to improve this manuscript for final publication in a peer-reviewed journal. Interested collaborators may contact the corresponding author.

## Completing Interest

The author declares no competing interest.

## Data Availability

The datasets generated during and/or analysed during the current study are available from the corresponding author on reasonable request. ITS, LSU and RPB2 sequences have been deposited in GenBank (Accession: PX630179).

## Ethics approval

This study does not involve human participants. Clinical bacterial isolates were obtained from State Specialist Hospital, Okitipupa with ethical approval No: SP/OK/2026/00023. This study was performed in line with the principles of the Declaration of Helsinki where applicable.

## Consent to participate

Not applicable as no human participants were involved.

## Consent to publish

Not applicable.

## Use of AI

Ai-assisted copy editing was used to improve readability and style. All content was human-generated and verified.

## References

Adedokun, O. M., Odiketa, J. K., Afieroho, O. E., & Afieroho, M. C. (2022). Importance of mushrooms for food security in Africa. In Sustainability sciences in Asia and Africa (pp. 343-360). Springer Nature Singapore. 10.1007/978-981-16-6771-8_20

Adetuwo, O. J., Fakoya, S., Ogundana, F. N. and Abeegunrin, T. A. (2026). Antifungal and bioactive potential of Pleurotus ostreatus cultivated on agro-waste substrates with molecular identification and functional characterization. Microbes & Immunity. doi: 10.36922/MI026160037

Ahmadi, A., Jamialahmadi, T., & Sahebkar, A. (2022). Polyphenols and atherosclerosis: A critical review of clinical effects on LDL oxidation. Pharmacological Research, 184, 106414. 10.1016/j.phrs.2022.106414

Ahmed, O. M., Ebaid, H., El-Nahass, E. S., Ragab, M., & Alhazza, I. M. (2020). Nephroprotective effect of Pleurotus ostreatus and Agaricus bisporus extracts and carvedilol on ethylene glycol-induced urolithiasis: Roles of NF-κB, p53, Bcl-2, Bax and Bak. Biomolecules, 10(9), 1317. 10.3390/biom10091317

AlAli, M., Alqubaisy, M., Aljaafari, M. N., AlAli, A. O., Baqais, L., Molouki, A., et al. (2021). Nutraceuticals: Transformation of conventional foods into health promoters/disease preventers and safety considerations. Molecules, 26(9), 2540. 10.3390/molecules26092540

Alberto, E. (2021). Naturally occurring strains of edible mushrooms: A source to improve the mushroom industry. In C. Z. Diego & A. Pardo-Giménez (Eds.), Edible and medicinal mushrooms (pp. 415–425). John Wiley & Sons. 10.1002/9781119149446.ch19

Ali, A., Ahmad, U., Akhtar, J., Badruddeen, J., & Khan, M. M. (2019). Engineered nanoscale formulation strategies to augment efficiency of nutraceuticals. Journal of Functional Foods, 62,103554. 10.1016/j.jff.2019.103554

Allam, E. A. H. (2024). Urolithiasis unveiled: Pathophysiology, stone dynamics, types, and inhibitory mechanisms: A review. African Journal of Urology, 30(1) 34. 10.1186/s12301-024-00436-z

Anachad, O., Taouil, A., Taha, W., & Bennis, F. (2023). The implication of short-chain fatty acids in obesity and diabetes. Microbiology Insights, 16, 1–10. 10.1177/11786361231162720

Andrade, G. M., Souza, E. L. D., Zárate-Salazar, J. R., et al. (2024). Unveiling the potential prebiotic effects of edible mushroom Pleurotus djamor during in vitro colonic fermentation. Journal of Agricultural and Food Chemistry, 72(48), 26722–26732. 10.1021/acs.jafc.4c06620

Aremu, M. O., Andrew, C., & Atere, A. (2018). Proximate and mineral composition of edible mushrooms. Journal of Food Measurement and Characterization, 12,2142–2151. 10.1007/s11694-018-9828-5

AOAC International. (2020). Official methods of analysis of AOAC International (21st ed.). AOAC International. 10.1093/9780197610145.001.0001

Bamisi, O. E., Ogidi, C. O., & Akinyele, B. J. (2024). Antimicrobial metabolites from probiotics, Pleurotus ostreatus and their co-cultures against foodborne pathogens isolated from ready-to-eat foods. Annals of Microbiology, 74, 31. 10.1186/s13213-024-01776-5

Barros, L., Cruz, T., Baptista, P., Estevinho, L. M., & Ferreira, I. C. F. R. (2008). Wild and commercial mushrooms as source of nutrients and nutraceuticals. Food and Chemical Toxicology, 46(8), 2742–2747. 10.1016/j.fct.2008.04.030

Benson, D. A., Cavanaugh, M., Clark, K., Karsch-Mizrachi, I., Lipman, D. J., Ostell, J., & Sayers, E. W. (2013). GenBank. Nucleic Acids Research, 41(D1), D36–D42. 10.1093/nar/gks1195

Benzie, I. F. F., & Strain, J. J. (1996). The ferric reducing ability of plasma (FRAP) as a measure of “antioxidant power: The FRAP assay. Analytical Biochemistry, 239(1), 70–76. 10.1006/abio.1996.0292

Boa, E. (2004). Wild edible fungi: A global overview of their use and importance to people. Food and Agriculture Organization of the United Nations.

Brand-Williams, W., Cuvelier, M. E., & Berset, C. (1995). Use of a free radical method to evaluate antioxidant activity. LWT - Food Science and Technology, 28(1), 25–30. 10.1016/S0023-6438(95)80008-5

Caglarirmak, N. (2007). The nutrients of exotic mushrooms (Lentinula edodes and Pleurotus species) and an estimated approach to the volatile compounds. Food Chemistry, 105(3), 1188–1194. 10.1016/j.foodchem.2007.01.004

Cai, M., & Hyde, K. D. (2019). Taxonomy of edible and medicinal mushrooms: Challenges and opportunities. Fungal Diversity, 97, 1–15. 10.1007/s13225-019-00427-9

Chang, S. T., & Miles, P. G. (2004). Mushrooms: Cultivation, nutritional value, medicinal effect, and environmental impact (2nd ed.). CRC Press. https://sayedmaulana.files.wordpress.com/2011/02/mushrooms.pdf

Chang, S. T., & Wasser, S. P. (2017). The role of culinary-medicinal mushrooms on human welfare with a pyramid model for human health. International Journal of Medicinal Mushrooms, 19(2), 95–134. 10.1615/IntJMedMushr.v19.i2.10

Chen, L., Gong, Y., Cai, Y., Liu, W., Zhou, Y., Xiao, Y.,…& Bian, Y. (2016). Genome sequence of the edible cultivated mushroom Lentinula edodes (Shiitake) reveals insights into lignocellulose degradation. PLOS ONE, 11 (8), e0160336. 10.1371/journal.pone.0160336

Clinical and Laboratory Standards Institute. (2018). Methods for dilution antimicrobial susceptibility tests for bacteria that grow aerobically (M07-Ed11). Clinical and Laboratory Standards Institute.

Effiong, M. E., Umeokwochi, C. P., & Afolabi, I. S. (2024). Comparative antioxidant activity and phytochemical content of five extracts of Pleurotus ostreatus. Scientific Reports, 14, 3794. 10.1038/s41598-024-54201-x

European Committee on Antimicrobial Susceptibility Testing. (2023). Antimicrobial susceptibility testing EUCAST guidelines. https://www.eucast.org/

Fakoya, S., Adegbehingbe, K. T., & Ademakinwa, I. S. (2020). Bio-therapeutic, phytochemical screening and antioxidant efficacies of oyster mushroom (Pleurotus ostreatus) obtained from the wild. Open Journal of Medical Microbiology, 10, 58–70. 10.4236/ojmm.2020.102006

Ferreira, I. C. F. R., Barros, L., & Abreu, R. M. V. (2009). Antioxidants in wild mushrooms. Current Medicinal Chemistry, 16(12), 1543–1560. 10.2174/092986709787909587

Ferraro, V., Gargano, M. L., Procida, G., Venturella, G., Cirlincione, F. & Cateni F. (2021). Mycochemicals in wild and cultivated mushrooms: nutrition and health. Phytochemistry Reviews, 21(2):339–383. doi: 10.1007/s11101-021-09748-2

Guzmán, G. (2008). Diversity and use of traditional medicinal fungi. International Journal of Medicinal Mushrooms, 10(3), 209–217. 10.1615/IntJMedMushr.v10.i3.20

Hasan, G. Q., & Abdulhadi, S. Y. (2022). Molecular characterization of wild Pleurotus ostreatus (MW457626) and evaluation of β—glucans polysaccharide activities. Karbala International Journal of Modern Science, 8(1), 52–62.10.33640/2405-609X.3204

Lavelli, V., Proserpio, C., Gallotti, F., Laureati, M., & Pagliarini, E. (2018). Circular reuse of bio-resources: The role of Pleurotus spp. in the development of functional foods. Food & Function, 9(3), 1353–1372. 10.1039/C7FO01747B

Mayirnao, H., Sharma, S. K., Jangir, P., Kaur, S., & Kapoor, R. (2024). Mushroom-derived nutraceuticals in the 21st century: An appraisal and future perspectives. Journal of Future Foods, 5(4), 342–360. 10.1016/j.jfutfo.2024.07.013

Newman, D. J. & Cragg, G. M.(2020). Natural products as sources of new drugs over nearly four decades from 01/1981 to 09/2019. J Nat Prod., 83(3):770–803. doi:10.1021/acs.jnatprod.9b01285

Oyetayo, V. O. (2012). Medicinal uses of mushrooms in Nigeria: towards full and sustainable exploitation. African Journal of Biotechnol., 10(12):2581–2593. doi:10.4314/ajtcam.v8i3.65289

Patel, S., & Goyal, A. (2012). Recent developments in mushrooms as anti-cancer therapeutics: A review. 3 Biotech, 2(1), 1–15. 10.1007/s13205-011-0036-2

Paterson, R. R. M. (2006). Ganoderma - A therapeutic fungal biofactory. Phytochemistry, 67(18), 1985–2001. 10.1016/j.phytochem.2006.07.004

Ragunathan, R., & Swaminathan, K. (2003). Nutritional status of Pleurotus spp. grown on various agro-wastes. Food Chemistry, 80(3), 371–375. 10.1016/S0308-8146(02)00275-3

Rathore, H., Prasad, S., & Sharma, S. (2017). Mushroom nutraceuticals for improved nutrition and better human health: A review. PharmaNutrition, 5(2), 35–46. 10.1016/j.phanu.2017.02.001

Re, R., Pellegrini, N., Proteggente, A., Pannala, A., Yang, M., & Rice-Evans, C. (1999). Antioxidant activity applying an improved ABTS radical cation decolorization assay. Free Radical. Biology and Medicine, 26(9-10), 1231–1237. 10.1016/S0891-5849(98)00315-3

Reis, F. S., Barros, L., Martins, A., & Ferreira, I. C. F. R. (2012). Chemical composition and nutritional value of the most widely appreciated cultivated mushrooms: An inter-species comparative study. Food and Chemical Toxicology, 50(2), 191–197. 10.1016/j.fct.2011.10.056

Reis, F. S., Martins, A., Barros, L., & Ferreira, I. C. F. R. (2012). Antioxidant properties and phenolic profile of the most widely appreciated cultivated mushrooms: A comparative study between in vivo and invitro samples. Food and Chemical Toxicology, 50(5), 1201–1207. 10.1016/j.fct.2012.02.013

Roupas, P., Keogh, J., Noakes, M., Margetts, C., & Taylor, P. (2012). The role of edible mushrooms in health: Evaluation of the evidence. Journal of Functional Foods, 4(4), 687–709. 10.1016/j.jff.2012.05.003

Sánchez, C. (2010). Cultivation of Pleurotus ostreatus and other edible mushrooms. Applied Microbiology and Biotechnology, 85(5), 1321–1337. 10.1007/s00253-009-2343-7

Schoch, C. L., Seifert, K. A., Huhndorf, S., Robert, V., Spouge, J. L., Levesque, C. A., … & Schindel, D. (2012). Nuclear ribosomal internal transcribed spacer (ITS) region as a universal DNA barcode marker for Fungi. Proceedings of the National Academy of Sciences, 109(16), 6241–6246. 10.1073/pnas.1117018109

Shirur, M., Barh, A., & Annepu, S. K. (2021). Sustainable production of edible and medicinal mushrooms: Implications on mushroom consumption. In V. K. Hebsale (Ed.), Climate change and resilient food systems (Vol. 1, pp. 315-346). Springer Singapore. 10.1007/978-981-33-4538-6_12

Silva, S. O., Costa, S. M. G., & Clemente, E. (2002). Chemical composition of Pleurotus pulmonarius (Fr.) Quél., substrates and residue after cultivation. Brazilian Archives of Biology and Technology, 45(4), 531–535.10.1590/S1516-89132002000600018

Singleton, V. L., Orthofer, R., & Lamuela-Raventós, R. M. (1999). Analysis of total phenols and other oxidation substrates and antioxidants by means of Folin-Ciocalteu reagent. Methods in Enzymology, 299, 152–178. 10.1016/S0076-6879(99)99017-1

Stamets, P. (2000). Growing gourmet and medicinal mushrooms (3rd ed.). Ten Speed Press.

Tamang, J. P., Watanabe, K., & Holzapfel, W. H. (2016). Diversity of microorganisms in global fermented foods and beverages. Frontiers in Microbiology, 7, 377. 10.3389/fmicb.2016.00377

Valverde, M. E., Hernández-Pérez, T., & Paredes-López, O. (2015). Edible mushrooms: Improving human health and promoting quality of life. International Journal of Microbiology, 2015, 376387. 10.1155/2015/376387

Vamanu, E. (2012). In vitro antimicrobial and antioxidant activities of ethanolic extracts of Pleurotus ostreatus. Annals of Microbiology, 62(2), 469–476. 10.1007/s13213-011-0348-9

Wasser, S. P. (2010). Medicinal mushroom science: History, current status, future trends, and unsolved problems. International Journal of Medicinal Mushrooms, 12(1), 1–16. 10.1615/IntJMedMushr.v12.i1.10

Wasser, S. P. (2011). Current findings, future trends, and unsolved problems in studies of medicinal mushrooms. Applied Microbiology and Biotechnology, 89(5), 1323–1332. 10.1007/s00253-010-3067-4

Wasser, S. P. (2014). Medicinal mushroom science: Current perspectives, advances, evidences and challenges. Biomedical Journal, 37(6), 345–356. 10.4103/2319-4170.138318

White, T. J., Bruns, T., Lee, S., & Taylor, J. (1990). Amplification and direct sequencing of fungal ribosomal RNA genes for phylogenetics. In M. A. Innis, D. H. Gelfand, J. J. Sninsky, & T. J. White (Eds.), PCR protocols: A guide to methods and applications (pp. 315–322). Academic Press.

Younis, A. M., Wu, F. S., & El Shikh, H. H. (2015). Antimicrobial activity of extracts of the oyster culinary medicinal mushroom Pleurotus ostreatus (Higher Basidiomycetes) and identification of a new antimicrobial compound. International Journal of Medicinal

Zaidman, B. Z., Yassin, M., Mahajna, J., & Wasser, S. P. (2005). Medicinal mushroom modulators of molecular targets as cancer therapeutics. Applied Microbiology and Biotechnology, 67(4), 453–468. 10.1007/s00253-004-1787-z

